# Identifying SNP associations and predicting disease risk from Genome-wide association studies using LassoNet

**DOI:** 10.1101/2021.08.29.458051

**Authors:** Hussain M. Sajwani, Samuel F. Feng

**Affiliations:** Department of Mathematics, Khalifa University of Science and Technology, Abu Dhabi, United Arab Emirates; Khalifa University Centre for Biotechnology, Khalifa University of Science and Technology, Abu Dhabi, United Arab Emirates

**Keywords:** Genome-wide association studies, significant threshold, association analysis, disease risk prediction, genetic risk

## Abstract

In this paper, we show that under certain conditions LassoNet [1] should outperform threshold of significance (P-value) methods for identifying multi-SNP disease associations and predicting disease risk using data from Genome Wide Association Studies. To demonstrate this, we built a genotype-phenotype simulation to comprehensively benchmark each method’s performance in variant selection and in predicting disease risk. Our results suggest that LassoNet, with its ease of implementation, should be added to the biomedical informatician’s toolkit. We release code to replicate our results.

## 1 Scientific Background

Genome-wide association studies (GWAS) have revolutionized the study of genetics and complex diseases, identifying over 50,000 associations between variants, typically single-nucleotide polymorphisms (SNPs), and complex traits and disease. These discoveries are then used to augment prediction for a variety of traits and diseases including body mass index, hair color, type 2 diabetes, and Alzheimer’s disease [3, 4]. The vast majority of these studies rely on identifying some threshold for significance for identifying SNP associations, with most GWAS publications using *p* = 5 × 10^*-*8^ [4]. Others take a more nuanced approach towards finding an appropriate threshold for identifying associations by making use of additional structure in the data [5]. Still, the performance of these methods for predicting disease risk is low [4], as the conventional False Discovery Rate and Bonferroni corrections are too conservative which results in low true positive rates [5]. Thus, there is a need for improvements in these methods for identifying SNPs and for predicting disease risk using GWAS data.

## 2 Materials and Methods

LassoNet is a recently developed feature selection method for neural networks that enables global feature selection via a convex sparsity penalty, resulting in improved interpretability and regression [1]. We propose using the feature selection afforded by LassoNet as an improvement over selecting variants according to *p*-values. In this section, we describe relevant details of these two approaches and a custom GWAS simulation framework we developed in order to benchmark and compare them. Code to replicate all of our methods, along with the results and figures in Section 3, is freely available online at https://github.com/HussainMSajwani/LNvsP.

### 2.1 Variant selection using P-values (“P-thresholding”)

Given genotype and case/control phenotype data, we followed a standard procedure and calculated *p*-values using a *t*-test for the odds ratio for association between genotype and case/control (e.g. PLINK, [2]). Calculations were performed using custom code and the `statsmodel` package for Python 3.7. Variants were sorted and selected according to significance. For example, in order to select *k* = 10 SNPs, P-thresholding chose the SNPs with the smallest 10 *p*-values.

### 2.2 Variant selection using LassoNet

LassoNet augments a deep neural network approach to classifying samples into cases and controls [1]. In brief, let NN_*W*_ : ℝ^*d*^ → ℝ be a dense feed-forward deep neural network with weights *W*. For any *x* ∈ ℝ^*d*^ and *θ* ∈ ℝ^*d*^ define the LassoNet as *f* : ℝ^*d*^ → ℝ such that

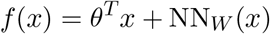

which can roughly be thought of as a residual neural network with one skip layer. Projected proximal gradient descent then repeatedly solves the following optimization problem:

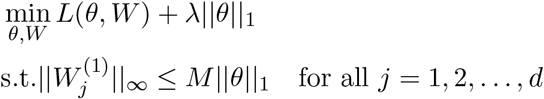

where ||*θ*||_1_ is the sum of the absolute values of the components of *θ* (the “Lasso” sparsity constraint). The LassoNet is trained many times with λ ← (1+ϵ)λ until all the weights from the first layer *W* ^(1)^ shrink to zero. This results in a regularization path of solutions, from which one can select any desired amount of feature sparsity. The importance of a feature (variant) *j* is determined by how far along the path (as λ increases) the weights 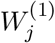zero out. Variants are selected by choosing the last *k* features to zero out in this way. The neural network used in all the following experiments has hidden layers of size 1200, 800, and 200, with the software default values of *M* = 10 and *ϵ* = 0.02.

### 2.3 Simulation benchmarks for comparing variant selection and disease prediction performance

Our GWAS simulations for comparing the performance between various methods begin with reference data (e.g. HapMap3, 1000 Genomes). A segment of *d* variants is extracted from a random location within some chosen chromosome, and then Hapgen2 [7] is used to simulate *n* genotype samples from this population data. These steps preserve the linkage disequilibrium (LD) structure from the original population, and are an important aspect of the disease prediction problem, even if several statistically associated SNPs are not truly causal.

Phenotypes were simulated using `PhenotypeSimulator` package in R [8]. This provided quantitative phenotypes, which were then labelled as cases and controls according to sign (+ or -). The phenotypes were generated in such a way that their variance is 95% explained by genotype and 5% explained by environmental factors. In other words, the phenotypes simulated in this study are highly heritable ones. The genetic contribution is then further broken down into two components. The first component is controlled by *d*_*c*_ causal SNPs and the second is controlled by the other *d − d*_*c*_ SNPs. In this paper, we always have *d*_*c*_ = 10 causal SNPs chosen uniformly at random. The effect size of the causal SNPs is set to be 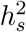 and the other SNPs have an effect size of 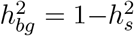. Thus 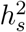 can be understood as controlling the polygenicity of the phenotype [6]. Any other parameter of `PhenotypeSimulator`’s `simulatePhenotypes` that was not discussed here was set to be the default values.

In order to estimate both model performance and variance, the process was repeated several times, each run producing new training data on which both methods were compared. The different methods indeed identified different associations, as illustrated in Figure 1. This simulation framework enables us to know “ground truth” about the causal SNP associations so that we can measure model performance in this domain. We also compare predictive performance for disease for each approach by taking their identified variants as inputs to either a logistic regression model or support vector machine (SVM). A 80/20 train/test split was used to measure the accuracy. Logistic regression was fit using the `statsmodels` version 0.12.0 package in Python. The `bfgs` algorithm is used to optimize the logistic regression model. The SVM was fit with `scikit-learn` version 0.23.2 with default parameters.

**Figure 1:**
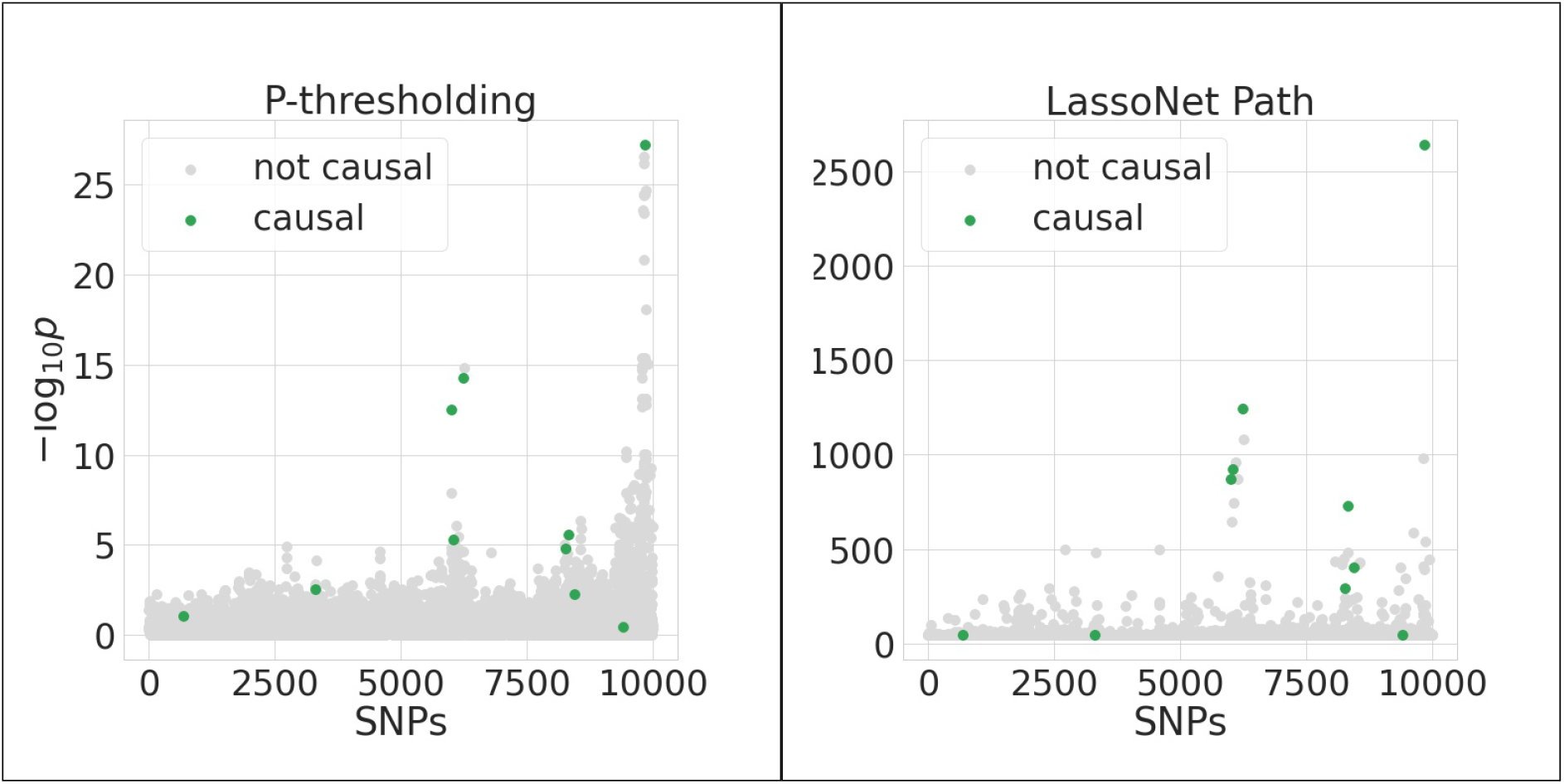
Each method attempting to identify 10 causal SNPs from simulated GWAS data.

## 3 Results

We carried out our study using genotype data from chromosome 22 of the CEU subpopulation from HapMap3 [10]. All results are reported over 50 independent simulations of the data. Phenotypes were generated using 10 causal SNPs. Each set of results contains comparisons between P-thresholding and LassoNet for a variety of values of *k* (*k* = 5, 25, 100, 300) and 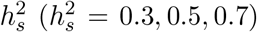. Recall from above that *k* is the number of variants selected and 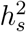 is the effect size of the causal SNPs. ∗, ∗∗, and ∗ ∗ ∗ denote significant differences in out of sample test performance at *p* < .05, .01, and .001, respectively.

Figure 2 shows each algorithms’ ability to correctly recover the 10 causal/associated SNPs for different values of *n, k*, and 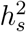. The left 12 subpanels show results for *n* = 480 samples, and the right 12 subpanels show results for *n* = 960. We observe that for both sample sizes, LassoNet correctly identifies significantly more causal

**Figure 2:**
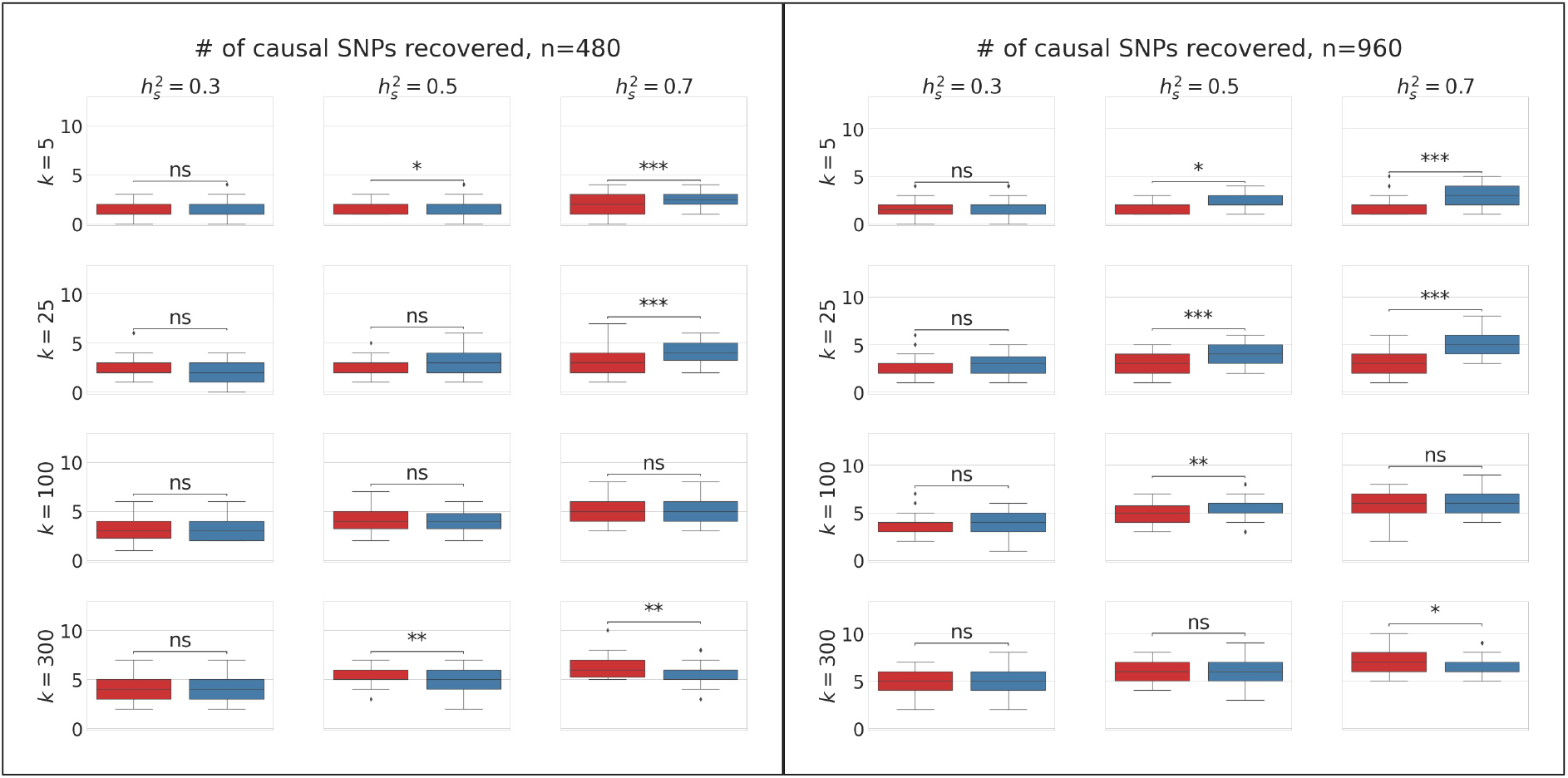
Variant selection performance of P-thresholding (red) vs. LassoNet (blue) for different sample sizes *n*. Left frame: *n* = 480. Right frame: *n* = 960.

SNPs compared to P-thresholding for moderate to high values of 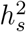and lower values of *k*. This roughly corresponds to diseases/phenotypes for which a moderate to large amount of variation can be explained by a specific subset of causal SNPs. We also observe that the improvements of LassoNet vs. P-thresholding are larger for *n* = 960 compared to *n* = 480, suggesting that LassoNet should also perform better even with larger sample sizes.

Figure 3 compares model performance across different numbers of non-causal SNPs, analogous to different sizes of SNP arrays collecting the data. Here we still

**Figure 3:**
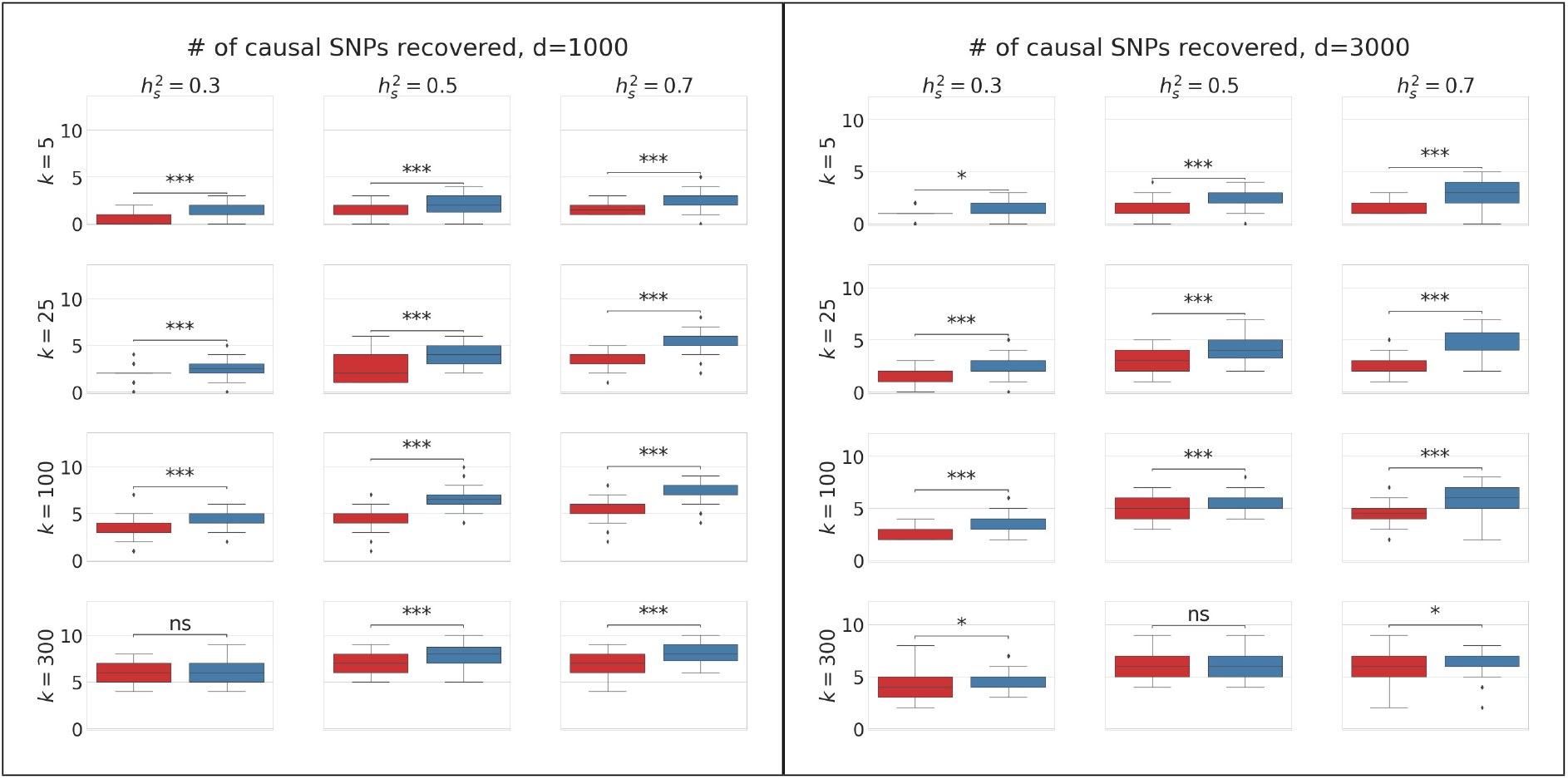
Variant selection performance of P-thresholding (red) vs. LassoNet (blue) for different numbers of measured SNPs (*d*). Left frame: *d* = 1000. Right frame: *d* = 3000

have 10 causal SNPs, fix *n* = 960, and change the total number of SNPs in the simulated data (*d* = 1000 or 3000). We again observe superior performance for LassoNet compared to P-thresholding over both sets of calculations.

Figure 4 shows the out of sample predictive performance for each method on an additional 240 independent test samples, assessing accuracy with the area under curve (AUC) criterion [9]. Each approach’s selected variants were used as inputs to a standard classification algorithm. The left 12 subpanels show results from fitting a logistic regression model with each method’s selected SNPs and the right 12 subpanels show results from using a support vector machine. Overall, we observe trends that LassoNet outperforms P-thresholding, especially when 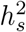is 0.5 or 0.7. We note that for disease risk prediction is strictly easier than the SNP identification problem above (Figures 2 and 3), as binary predictions with good accuracy can be made without correctly identifying all causal SNPs.

**Figure 4:**
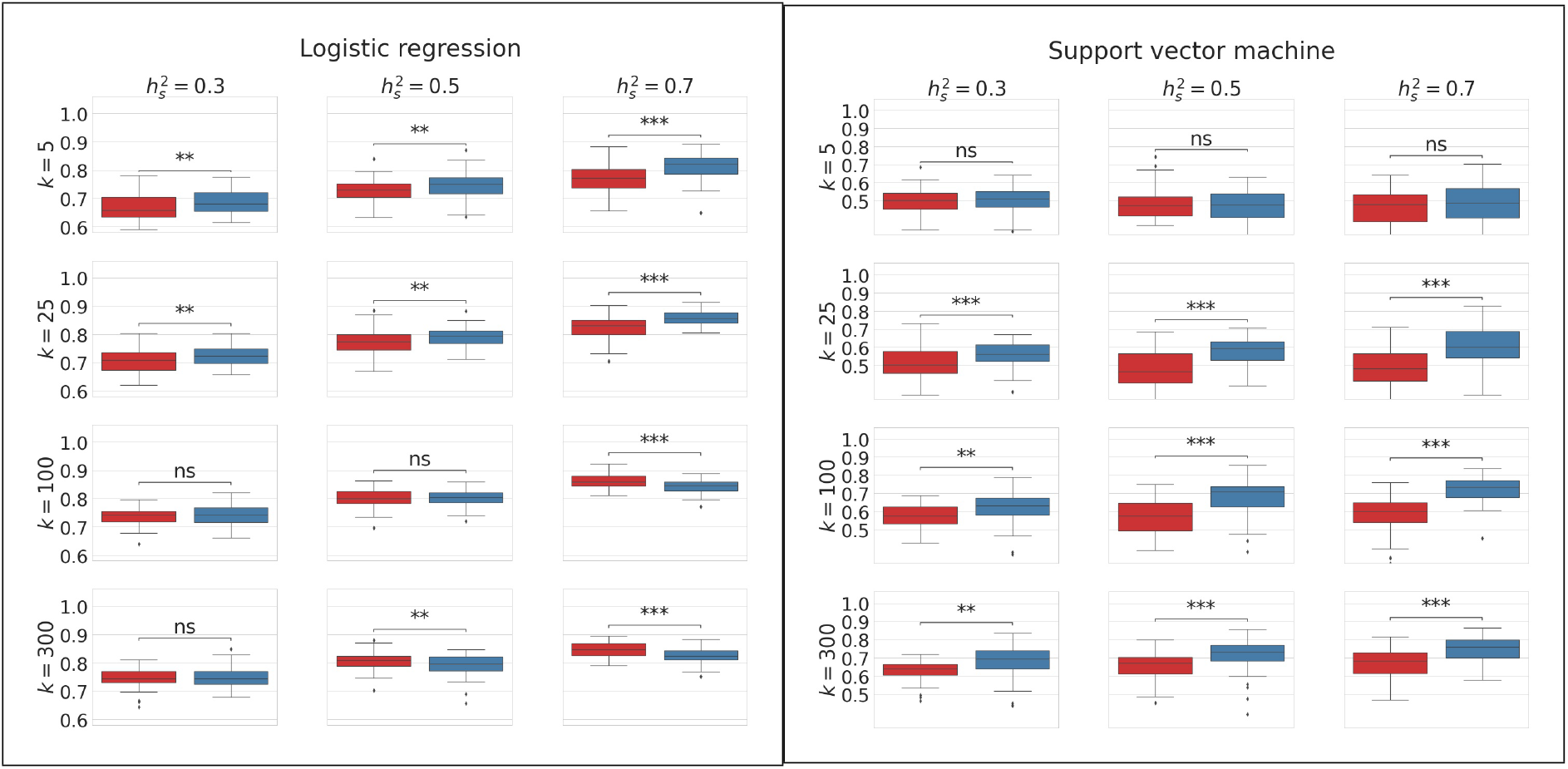
Disease risk prediction performance (AUC) of P-thresholding (red) vs. LassoNet (blue). Left frame: Prediction via logistic regression. Right frame: Prediction via support vector machine.

All simulations were performed on a system running Ubuntu 18.04.5 LTS, 12 Xeon E5-2603 cores and 64GB RAM. Training a LassoNet with *n* = 1200 and *d* = 10000 with the architecture stated above took an average of 24.5GB of RAM (stdev = 3.5GB) and an average of 16.0 minutes (stdev = 10 minutes.)

We note that our GWAS simulation pipeline enables generating similar robust results for a variety of other conditions such as different amounts of environmental effects, polygenic effects, quantitative traits, and population substructure. Furthermore, one may easily compare different prediction and inference algorithms to P-thresholding and LassoNet.

## 4 Conclusion

We have demonstrated that LassoNet [1] outperforms P-thresholding methods both in identifying SNPs associated with disease as well as in predicting disease risk using GWAS data. Our simulations suggest that these advantages are observed for moderate to high levels of genetic effect size and over different sample sizes, numbers of SNPs filtered, and different sizes of SNP arrays. This means that LassoNet captures more causal/associated SNPs than P-thresholding, in addition to maintaining higher overall accuracy (AUC) for predicting disease when coupled with logistic regression or support vector machines. LassoNet’s superior performance was more robust for more severe filtering criteria (i.e. forced to select fewer variants). These results suggest that LassoNet should be part of the medical informatician’s toolkit for selecting SNPs and predicting disease risk. We have released code to replicate our findings.

## Acknowledgments

This paper is based upon work supported by the Khalifa University of Science and Technology under Award No. CIRA-2019-050 to SFF.

